# CRISPR-Cas9 Screen Reveals PSMB3 Contributes to Gliomagenesis Through Proteasome-Dependent and Independent Mechanisms

**DOI:** 10.1101/2021.07.28.453566

**Authors:** Shivani Baisiwala, Shreya Budhiraja, Andrew J Zolp, Khizar Nandoliya, Li Chen, Cheol H. Park, Ella N Perrault, Miranda R Saathoff, Crismita Dmello, Jack M Shireman, Peiyu Lin, Gabriel Dara, Katy McCortney, Craig Horbinski, Adam M Sonabend, Atique U Ahmed

**Affiliations:** Department of Neurological Surgery, Feinberg School of Medicine, Northwestern University, Chicago, IL 60611, USA; Northwestern University, Evanston, IL, 60208

**Author notes:** <u>Correspondence should be addressed to:</u> Atique U. Ahmed, PhD, Department of Neurological Surgery, Feinberg School of Medicine, Northwestern University,303 East Superior Street Chicago, IL60611. Shivani Baisiwala, MD MS, Department of Neurological Surgery, Feinberg School of Medicine, Northwestern University,303 East Superior Street Chicago, IL60611.

## Abstract

Glioblastoma (GBM) is the most common adult malignant brain tumor, with a median survival of 21 months and a 100% recurrence rate. Even though many of the critical oncogenic drivers for GBM have been identified, the basis of gliomagenesis is still under investigation. To identify novel genes that contribute to GBM progression, we performed a genome-wide CRISPR-Cas9 knockout screen. We identified four previously unstudied genes – PSMB3, CHCHD4, SPDYE5, HSPA1 – which had elevated expression in cancer and demonstrated a significant positive correlation with respect to GBM growth and patient survival in vivo and patient datasets. Furthermore, overexpression of PSMB3 and HSPA5 in neural stem cells resulted in transformation to a cancer phenotype. Further investigation of PSMB3, a subunit of the proteasome, allowed us to identify both ubiquitin-mediated and non-ubiquitin-mediated mechanisms of oncogenesis. Ultimately, the data from our CRISPR screens suggests that these genes drive tumor progression, making them promising therapeutic targets for GBM.

## Introduction

Glioblastoma (GBM) is the most common adult primary brain malignancy. Even with an aggressive standard of care that combines maximal surgical resection with radiation and temozolomide (TMZ)-based chemotherapy, GBM remains an extremely aggressive tumor with universal recurrence in patients and a median survival of 21 months [1–6]. This inevitable recurrence is thought to be due to the vast heterogeneity of the tumor, described previously by many holistic genetic studies [5, 6]. Despite years of research, GBM remains incurable, with little progress made over the past few decades [3]. Thus, there is an urgent need to identify novel druggable targets and mechanisms. One approach to identification of these targets is through unbiased functional screening to identify previously unstudied targets.

CRISPR (clustered regularly interspaced short palindromic repeats)-Cas9 is a novel technology that introduces highly directed genomic edits in an efficient and accurate manner [7]. Recent developments in CRISPR-Cas9 technology allow it to be used in a high-throughput fashion in order to perform genome-wide screens [8]. This approach allows for a unbiased screen across large sets of genes to determine which ones directly and functionally contribute to a phenotype of interest [9]. Thus, CRISPR-Cas9 screening enabled us to identify novel targets that drive GBM’s fitness and progression.

Here we show that conducting a genome-wide screen in glioma cells resulted in the depletion of nearly 160 novel essential growth genes. We further identified critical pathways contributing to GBM’s fitness. Four previously understudied genes in GBM - PSMB3, CHCHD4, SPDYE5, HSPA5 - were chosen for this study based on novelty, depletion within our screen, and clinical relevance. We further establish a mechanism for PSMB3 as a key component of the ubiquitin-proteasome system (UPS), which is a multi-catalytic complex that recycles ubiquitinated intracellular proteins. The UPS helps maintain protein homeostasis within the cell and thus has been associated as a target for cancer phenotypes and therapies in multiple previous studies [10–13] [14–16]. In addition, we identify an additional mechanism of PSMB3 as an independent regulator of Notch activity, which is a known critical driver for glioblastoma. The role of PSMB3 in tumorigenesis remains unstudied in GBM. Our goal here was to understand genes and pathways contributing to gliomagenesis and then to elucidate the role of PSMB3 in gliomagenesis both within and outside the UPS.

## Materials & Methods

### CRISPR-Cas9 Knockout Screening

Screening was performed as previously described [17]. Briefly, H4 human GBM cells were infected with the whole-genome knockout Brunello library (Addgene, Cambridge, MA, USA), which covered ~19,000 genes with 4 sgRNAs per gene along with 10,000 sgRNA non-targeting controls. In order to prepare the library, we harvested and seeded 80% confluent HEK293T cells into a T225 flask for 20-24 hours. Opti-MEM I reduced serum, pMD2.G −5.2 µg/ml, psPAX −10.4 µg/ml, and Lipofectamine plus reagent was then added to the cells. After the cells were incubated for 4 h, 0.45 µM filters were used to filter the media, whereafter the virus was then aliquoted and stored at −80 °C.

To titer the virus, 2 ml of media and 3 million cells were seeded into a 12-well plate. Cells were transduced with either 400, 200, 100, 75, 50, 25, or 0 μl of virus, with 8 µg/µl of polybrene in each well. Cells were then spinfected at 1000 g at 33 °C for 2 h. After incubating them for 24 hours at 37 °C, cells were harvested and seeded at a density of 4000 cells for 96 h. A non-transduced cell control was also maintained in similar conditions. All cells were treated with puromycin and then a cell titer glo assay was used to determine viability and calculate the multiplicity of infection (MOI).

Once titer was determined, 500 million H4 cells were cultured and spinfected with the sgRNA library. After 4 days selection with 0.6 µg/ml of puromycin, approximately 150 million cells remained. Of these cells, a subset was used to extract genomic DNA (gDNA) after amplification of sgRNA with unique barcoded primers, serving as the day 0 control. The rest of the cells were expanded for 14 days, after which another subset was extracted, and then 28 days, at which point the final set was extracted.

In order to amplify the sgRNA, the gDNA was extracted using the Zymo Research Quick-DNA Midiprep Plus Kit (Cat no: D4075, Irvine, CA, USA). After gDNA was cleaned with 100% ethanol with 1/10 volume 3 M sodium acetate, PH 5.2, and 1:40 glycogen co-precipitant (Invitrogen Cat no: AM9515), Nano drop 2000 (Thermo Scientific, Waltham, MA, USA) was used to measure gDNA concentration. PCR was then performed to expand the library and next generation sequencing (Next Seq) was used to sequence the sgRNAs at 300 million reads for the four sgRNAs pool aiming at 1000 reads/sgRNA. The samples were sequenced with 80 cycles of read 1 (forward) and 8 cycles of index 1, as stated in the Illumina protocol. In order to enhance coverage, 20% PhiX was added on the Next Seq.

Analysis of computational data was performed using CRISPRAnalyzR and the CaRpools pipeline. Both the DESeq2 and the MaGeCK algorithms were applied to determine significance [18].

### Enrichment Mapping

Significant gene lists were first identified from CRISPR or mass spectrometry per appropriate analysis methods. Lists were entered into gProfiler and the GEM file was downloaded. The EnrichmentMap app within Cytoscape was used to load the enrichment map and group pathways appropriately [19–21].

### Cell Lines & Culture

U251, a human glioma cell line, was procured from the American Type Culture Collection (Manassas, VA, USA). Dulbecco’s Modified Eagle’s Medium (DMEM; HyClone, Thermo Fisher Scientific, San Jose, CA, USA), with 10% fetal bovine serum (FBS; Atlanta Biologicals, Lawrenceville, GA, USA) and 1% penicillin-streptomycin antibiotic mixture (Cellgro, Herndon, VA, USA; Mediatech, Herndon, VA, USA), were used to culture the cells. The patient-derived xenograft (PDX) glioma cells (GBM43, GBM5, and GBM6) were procured from Dr. C. David James at Northwestern University and maintained according to published protocols [1, 2]. These PDX cells were cultured in DMEM, with 1% FBS and 1% penicillin-streptomycin. A frozen stock was used to replenish cells that were already used for a maximum of 4 passages. Frozen cells were maintained in liquid nitrogen at −180°C in pure FBS supplemented with 10% dimethyl sulfoxide (DMSO).

Neural stem cell, fibroblast, and astrocyte lines were obtained from American Type Culture Collection (Manassas, VA, USA) or from Kerafast (Boston, MA, USA). H1B.F3 lines and fibroblast (WI-38) lines were cultured in Dulbecco’s Modified Eagle’s Medium (DMEM; HyClone, Thermo Fisher Scientific, San Jose, CA, USA), with 10% fetal bovine serum (FBS; Atlanta Biologicals, Lawrenceville, GA, USA) and 1% penicillin-streptomycin antibiotic mixture (Cellgro, Herndon, VA, USA; Mediatech, Herndon, VA, USA). LM NSC008 cells were cultured in neurobasal media (Thermo Fisher Scientific, San Jose, CA, USA) supplemented with B27 (no Vitamin A; Invitrogen, Carlsbad, CA, USA), basic fibroblast growth factor (bFGF; 10 ng/mL; Invitrogen), epidermal growth factor (EGF; 10 ng/mL; Invitrogen), and N2 (Invitrogen)., and 1% penicillin-streptomycin solution (Cellgro). Media was replaced every 2 days for optimal growth. Astrocytes were cultured in astrocyte media (Thermo Fisher Scientific, San Jose, CA, USA) supplemented with N2 (Invitrogen), 10% fetal bovine serum (FBS: Atlanta Biologicals, Lawrenceville, GA, USA) and 1% penicillin-streptomycin solution (Cellgro).

### Cellular Transfection

Low passage 293T cells (ATCC, Manassas, VA, USA) were plated at 90% confluency in order to generate lentiviral particles. After 6 hours, the cells were then transfected with a mix of HP DNA Transfection Reagent (Sigma Aldrich, St Louis, MO, USA) diluted in OptiMEM media (Gibco, Waltham, MA, USA) as well as packaging and target plasmids, in accordance with manufacturer’s instructions. CRISPR-Cas9 knockout plasmids were obtained from Genecopoeia (Rockville, MD, USA) and overexpression plasmids were obtained from GenScript (Piscataway, NJ, USA). Other cell lines transfected for non-viral purposes were transfected in a similar fashion, with amounts scaled to dish area. Transfections were maintained for 48-72 hours before assaying transfection success.

### Viral Transduction

After the transfected cells were maintained in culture for 48-72 hours, the supernatant containing the virus was harvested, centrifuged at 1200 rpm for 5 minutes, and sterilized with a 45-micron filter. Viral supernatant was then ultracentrifuged for 2 hours at 4C to isolate viral particles. Particles were resuspended in PBS and maintained in the –80 until use. When ready for transduction, cells were resuspended in ~50 ul of media, and ~20 MOI lentivirus amounts were added per sample and 1mL of appropriate media. The virus-media mixture was spun for 2 hours at 37 °C at 850g. Following centrifugation, cells were plated and maintained in culture in regular media in addition to the viral media they were centrifuged in. After 2-3 days, media was changed to non-viral-containing media and cells were assayed for efficiency by western blot. After incubating overnight at room temperature, the cells were plated and maintained in culture with regular media changes for 48-72 hours.

### Animals & In Vivo Models

Athymic nude mice (nu/nu; Charles River, Skokie, IL, USA) were used in this study and were housed in compliance with Institutional Animal Care and Use Committee (IACUC) requirements along with federal and state statutes. Animals were subject to a 12-hour light and dark cycle with food and water available in the shoebox cages.

Intracranial implantation of glioblastoma cells was performed according to our lab’s previously established glioblastoma mouse model [1]. In this model, animals first received buprenex and metacam by intraperitoneal (IP) injection. They then were anesthetized from a second injection of ketamine/xylazine mixture (Henry Schein; New York, NY, USA). Mice were pinched in the foot to confirm complete sedation. Then, artificial tears were then applied to each eye, and ethanol and betadine were applied to the scalp for sterilization. The skull was then exposed by creating a small incision using a scalpel, whereafter a ~1mm burr hole was drilled right above the right frontal lobe. Then, the mice were placed in a stereotactic rig, where 5 μl of cell solution loaded in a Hamilton syringe was injected 3 mm from the dura over a period of one minute. In order to ensure that the cell suspension was released, the needle was then raised slightly and left for an additional minute. The syringe was then slowly removed, and the scalp was closed with sutures (Ethicon; Cincinnati, OH, USA). The position of the head was maintained in the same place throughout the closing process. Following surgery, animals were placed on heat pads until awake and reactive.

Throughout the study, signs of tumor progression -- including weight reduction, reduced body temperature, and hunching -- were monitored for and recorded. After determining that animals were unlikely to survive the next morning, animals were euthanized according to Northwestern University and IACUC guidelines and brains were cryopreserved for further assays.

### Immunofluorescence

Mouse brains were harvested and frozen in cryoprotectant on dry ice, stored at −80 °C, sectioned at 8 *μ*m per section, and stained according to standard immunohistochemistry protocols. In short, sections were thawed at room temperature for 15-20 minutes. They were then washed 2 times for 5 minutes each in PBS + 0.05% tween (PBS-T) to remove any remaining cryoprotectant. Next, the immunopen was used to draw a circle around each brain section. Sections were fixed in ~100ul of 4% PFA at room temperature for 15 minutes and then washed 2 times for 5 minutes each. Subsequently, antigen retrieval was performed by boiling samples in sodium citrate buffer for approximately 20 minutes, followed by a 30 minute cooling period to allow slides to return to room temperature. Slides were then washed 3 times for 5 minutes in PBS-T and then blocked and permeabilized in a 10% BSA solution with Triton-X (Thermo Fisher Scientific, Rockford, IL, USA) for 2 hours at room temperature. Then, sections were incubated overnight at 4 °C with ~100ul primary antibodies diluted in 1% BSA+Triton-X (Thermo Fisher Scientific, Rockford, IL, USA) [1]. The next morning, sections were washed 3 times for 10 minutes each in PBS-T. Approximately 100ul of secondary antibodies diluted in 1% BSA+Triton-X were added (Thermo Fisher Scientific, Rockford, IL, USA). After incubation for 2.5 hours at room temperature, sections were washed in PBS-T 3 times for 10 minutes each. Lastly, a drop of prolong gold anti-fade reagent with DAPi was added to each section (Thermo Fisher Scientific, Rockford, IL, USA), and slides were mounted and sealed with nail polish. Finally, Slides were imaged using a Leica confocal microscope [1]. Images were compiled and analyzed in ImageJ.

Additional experiments were conducted using immunocytochemistry protocol. Plates were first removed from the 37 °C incubator and washed once with PBS-T. Next, 200ul of 4% PFA was added to each well for 10 minutes at room temperature. Cells were then gently washed with PBS-T and subsequently blocked for 2 hours at room temperature in 200ul of 10% BSA solution. After blocking, the BSA was aspirated off of the slides, and 100ul of primary antibody mixed with 1% BSA were added. Cells were incubated at 4 °C overnight. Subsequently, cells were washed 3 times for 10 minutes each in PBS-T and 200ul of secondary antibody solution was added to each well for 2 hours are room temperature. Cells were then washed again 3 times for 10 minutes each in PBS-T. Phalloidin was then added at a concentration of 1:3000 to each well for 30 minutes at room temperature followed again by three 10 minute washes in PBS. A drop of prolong gold anti-fade reagent with DAPi was added to each section (Thermo Fisher Scientific, Rockford, IL, USA), and slides were mounted and sealed with nail polish. Finally, Slides were imaged using a Leica confocal microscope [1]. Images were compiled and analyzed in ImageJ.

### Cell Viability Assays

Viability assays were conducted using the MTT assay. Briefly, cells were plated at 3000-5000 per well in a 96-well plate with 6-8 replicates per condition. After the appropriate time period, the media was removed and cells were treated with MTT solution. This MTT solution was made by first diluting MTT stock reagent at 5mg/ml in dPBS. Next, this MTT stock was diluted in fresh media at a ratio of 1:10. 110ul of this mixture was added to each well, and cells were incubated at 37 °C for 4 hours. After incubation, the media was carefully removed and 100ul of DMSO was added to each well. Crystals were resuspended in the DMSO and the plate was incubated at room temperature for 10 minutes. Results were read out at an absorbance of 570nm and analysis was performed in excel to determine the percent viability relative to control.

### Western Blotting

Cells were trypsinized, washed in PBS, and resuspended in mammalian protein extraction reagent (M-PER; Thermo Scientific, Rockford, IL, USA) supplemented with protease and phosphatase inhibitor cocktail (PPI; Thermo Scientific, Rockford, IL, USA) and EDTA (Thermo Scientific, Rockford, IL, USA). Cells were then sonicated in a water bath for three 30 second increments, with 30 seconds of rest between each sonication. The resulting lysates were centrifuged at 13000 rpm for 10 min at 4 °C. Following supernatant collection, the protein concentration for each western blot sample was determined via BSA assay (Thermo Scientific, Rockford, IL, USA). Samples were then equalized and 6x sodium dodecyl sulfate buffer (SDS sample buffer; Alfa Aesar, Wood Hill, MA, USA was added. Samples were then boiled at 95 °C for 10 minutes in preparation for loading. Samples were then loaded and run through SDS-polyacrylamide gels made in-house at 45V initially and then 100V until completion. Proteins were transferred onto polyvinylidene difluoride (PVDF) membranes (Millipore, Darmstadt, Germany) at 14V constant in a semi-dry transfer machine. Subsequently, the membranes were rinsed in distilled water briefly and transferred to blocking solution (5% powdered milk in TBST) for one hour at room temperature. Membranes were then incubated in primary antibody solution overnight at 4°C. Primaries were made in 5% BSA solution supplemented with sodium azide. The next morning, membranes were washed in TBST 3 times for 10 minutes each and incubated in secondary antibody diluted 1:4000 in 5% milk. After washing the membranes 3 times for 10 minutes each, they were coated with enhanced chemiluminescence (ECL; Clarity ECL, BioRad), and images were developed a developer machine (BioRad, Hercules, CA, USA).

### Soft Agar Assay

Soft agar stock solution was made by adding 0.9g of agarose to 30ml of distilled water, after which the mixture was microwaved in 15 second intervals until the agarose completely dissolved. This solution was autoclaved for 15 minutes. 5ml and 10ml pipettes were warmed in the 37 °C incubator to prevent the agarose solution from solidifying while pipetting. 5ml of the agarose solution and 20ml of warmed media were transferred to a 50ml tube. After inverting the tube to mix, 2ml of this mixture was added to a 6-well plate and incubated for one hour to solidify.

Cells of interest were trypsinized and counted. Cell suspensions were made with 2.5mL of media and 25,000 cells. Once again, 5mL of the agarose solution and 20mL of warmed media were transferred to a 50mL tube to form the base agarose solution. 2.5mL of the base solution was added to 2.5mL of the cell suspension, mix briefly, and then distributed among 3 wells of a 6-well plate. This plate was then incubated for 1 hour to solidify and then 10 minutes at 4°C. Cells were then incubated at 37°C and 5% CO2, with feeding every 2-3 days, until colony formation was observed (approximately 3-5 weeks). Colonies were stained using crystal violet and counted in ImageJ.

### Flow Cytometry & Notch Reporter Assays

For notch reporter experiments, cells were previously transfected with the appropriate reporter. After 48-72 hours, cells were harvested and washed once with PBS. They were then collected in 80 μl of PBS and transported to the flow cytometry core in darkness. Samples were run on a Fortessa Analyzer and gated for live cells, single cells, and notch reporter positivity. Analysis was performed in FlowJo.

### Immunoprecipitation

Protein samples were isolated and normalized as above. Following normalization, 100ug of protein was incubated with 6 uL of ubiquitin antibody and protease/phosphatase inhibitors (Cell Signaling Technologies, Danvers, MA, USA) overnight. Subsequently, 30 uL of Protein A/G beads were added to the sample and it was incubated for 2 hours at room temperature. Reactions were then washed three times with 500 µl of MPER and spun down at 3200 g for 5 minutes. After the third wash, 50 µl of supernatant was kept and 50 µl of 2x SDS was added to all reactions. Samples were incubated at 55 C for 10 minutes to elute and then samples were spun down at 3200 g for 5 minutes to separate beads from the supernatant. The supernatant was collected and boiled at 95 C for 10 minutes and then directly loaded onto an SDS-PAGE gel. The remainder of the protocol was conducted per the western blotting protocol above or samples were sent for mass spectrometry with the Northwestern protein core. Analysis of mass spectrometry was performed using existing Scaffold Proteome Software (v4.11, Portland, OR, USA) and inbuilt statistics.

### Enrichment Mapping

Enrichment mapping was performed using gProfiler and Cytoscape. Briefly, genes of interest were inputted into gProfiler and appropriate enrichment functional pathways were selected (e.g. WikiPathways, etc.). The outputted GEM file was then uploaded into Cytoscape using the EnrichmentMap extension. This generated a base EnrichmentMap with pathways highlighted. Pathways were grouped and represented as a figure.

### Single Cell RNA-Seq

Single cell Drop-Seq was performed per previously published protocols [22]. Briefly, cells were isolated and formed into a single cell suspension. They were run through a Drop-Seq set up per established protocols and then sent out for sequencing. Analysis was performed using the Seurat pipeline [23].

Single cell data on localization was obtained from the freely available data browser at gbmseq.org [24].

### Proteasome Activity Assay

The proteasome activity assay was performed using a kit obtained from Abcam (Cambridge, UK). Briefly, cells were harvested, and lysate was isolated following the manufacturer’s protocol. A dark 96-well plate was set up with 5uL of lysate and the appropriate buffers in the kit. As soon as the reaction was started, the plate was read out on a plate reader continuously every 5 minutes.

### Quantitative PCR

RNA extraction was performed using Qiagen RNEasy kits (Qiagen, Hilden, Germany) per published protocols. The cDNA was generated from the RNA samples using an iScript kit (BioRad, Hercules, CA, USA) per manufacturer’s protocol. The cDNA was then diluted 1:10 in distilled water to use for downstream quantitative PCR. Reactions were set up with standard amounts of cDNA, SyberGreen (Thermo Fisher, Rockford, IL, USA), forward and reverse primers (IDT, Newark, NJ, USA) and read out on a standard qPCR machine. All primers were generated from Primer-BLAST using the fixed settings. Reactions were performed in triplicates.

### Neurosphere Assays

Cells were harvested, washed with PBS, and plated in serial dilutions, each with 12 replicates in neurobasal media (Gibco) supplemented with B27 (no Vitamin A; Invitrogen, Carlsbad, CA, USA), basic fibroblast growth factor (bFGF; 10 ng/mL; Invitrogen), epidermal growth factor (EGF; 10 ng/mL; Invitrogen), and N2 (Invitrogen). A blinded experimenter analyzed the wells after approximately 1 week. The number of neurospheres with a diameter greater than 20 cells was counted. Counts were analyzed using the Walter + Eliza Hall Institute of Medical Research platform (http://bioinf.wehi.edu.au/software/elda/). In addition, the absolute number of spheres was plotted visually, and images were taken of the wells using a Leica confocal microscope (Buffalo Grove, IL, USA).

### Statistical Analysis

Statistical analyses were performed using the GraphPad Prism v8.0 software (GraphPad Software; San Diego, CA, USA). In general, data are presented as mean with standard deviation for continuous variables and number or percentage for categorical variables. Differences between two groups were assessed using Student’s *t* test or Wilcoxon rank sum test where appropriate. Differences among multiple groups were assessed using ANOVA with *post hoc* Tukey’s test, or Mann–Whitney *U* test followed by Bonferroni correction where appropriate. Survival curves were graphed with the Kaplan-Meier method and compared by log-rank test. All tests were two-sided and p < 0.05 was considered statistically significant. Experiments such as Western blots, microarray, and fluorescence activated cell sorting analysis (FACS) were performed in biological triplicate. All *in vivo* experiments contained at least five animals per group. Males and females were equally represented, and each animal was treated as a technical replicate.

## Results

### Whole-Genome CRISPR Screen Identifies Essential Genes in GBM

To investigate which genes drive viability in GBM, we performed an unbiased whole-genome CRISPR-Cas9 knockout screen in H4 human GBM cells. This screen was performed using the whole-genome knockout Brunello library, which includes over 19,000 genes, including non-targeting controls, with coverage of 4 sgRNAs per gene. The screening was performed on days 14 and 28 to investigate two-time points (Figure 1A). Genomic sequencing quality assessments showed excellent per base sequence quality, appropriate quality score distribution over the sequence length, and the expected length distribution over all sequences (Figure S1A). Furthermore, the cumulative frequency of reads reflected similar distributions across conditions and our similarity analysis showed that samples were appropriately similar to replicates and varied across experimental conditions (Figure S1B, C). Overall results showed significant populations of depleted and enriched guides at both time points, with depleted guides corresponding to essential or driver genes and enriched guides to suppressor genes (Figure 1B). As a critical positive control, we investigated read counts of known crucial genes at day 0 versus day 28. We found that known essential genes were significantly depleted, reflecting the reliability of our screen results (Figure 1C). We further investigated specific genes found to be enriched and depleted in the screen. We were able to identify known tumor suppressors including TP53, BAX, and ATM, at both time points (Figure 1D). Our goal in this study was to identify those genes that drive oncogenic progression, and thus, we focus exclusively on the depleted guides or driver genes moving forward. Analysis showed that 222 genes were essential genes in our screen, of which only 86 are known crucial genes, and the remaining 136 are novel and understudied genes unique to GBM (Figure 1E). These essential genes were identified using multiple statistical analysis methods including DESeq2 and MAGeCK and multiple time points, thus reinforcing their importance in GBM (Figure 1F). In addition to specific genes, we performed enrichment analysis of our results to determine which pathways were most important in these processes. Analysis of suppressor pathways showed that DNA-damage response pathways, cancer signaling, and pro-apoptotic immune pathways were enriched (Figure 1G). In contrast, analysis of essential gene pathways reflected DNA repair processes, the proteasome, glycolysis, replication, and metabolism (Figure 1H).

**Figure 1:**
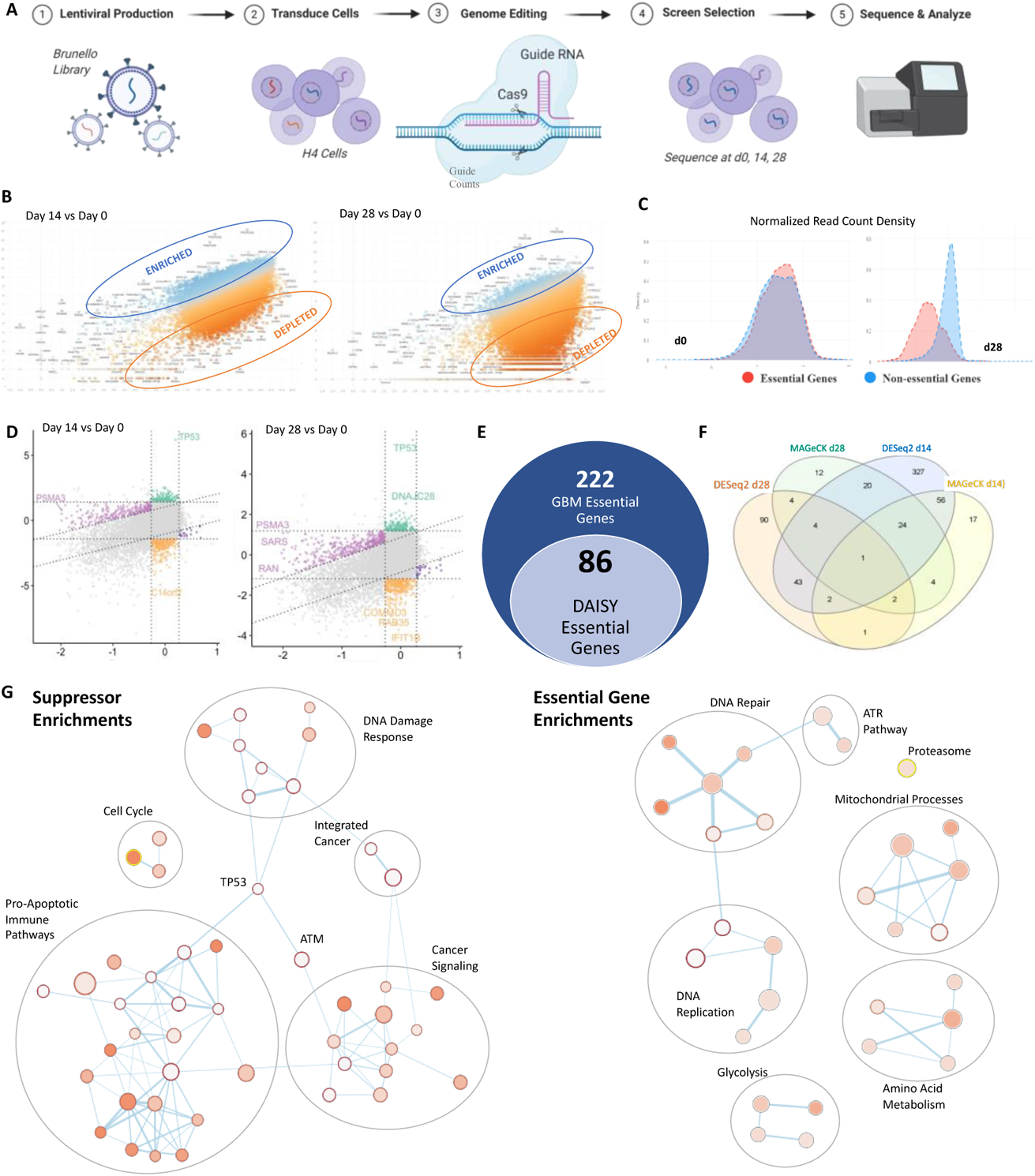
Whole-Genome CRISPR-Cas9 Knockout Screen Reveals Key Vulnerabilities in GBM. (A) CRISPR-Cas9 knockout screening was performed using the Brunello whole-genome library, with coverage of 19,000 guides including 4 guides per gene and ~2000 non-targeting controls. Sequencing was performed at days 0, 14, 28 (B) Comparison of day 14 and d28 versus day 0 revealed segments of guides that were enriched an depleted (C) Known essential gene read count densities were computed at day 0 and day 28. Essential genes are depleted at day 28, reflecting the validity of our screen (D) Specific genes were examined for enrichment and depletion at day 0, day 14, and day 28. Key suppressors were identified as enriched (BAX, TP53, ATM) at each timepoint, reflecting validity of the screen (E) 222 total genes were significantly depleted of which only 86 were previously known essential genes (F) Essential genes identified and pursued were identified across multiple timepoints with multiple CRISPR-appropriate analysis algorithms (G) Enrichment mapping identified key pathways across identified suppressors and drivers. Analysis was performed using the CaRPools pipeline, applying DESeq2 and MAGeCK algoirthms to identify significance. Enrichment mapping was performed in Cytoscape using the EnrichmentMap application.

### Identified Targets Are Upregulated in GBM and Impact Patient Survival

Our next goal was to experimentally validate targets of interest, having identified a broad list of genes and pathways implicated in the progression of glioblastoma. Initial analysis identified approximately 6000 significantly depleted guides. Of these, 222 were significant based on multiple analysis algorithms, and 17 met our criteria of being novel, clinically significant. From these, 4 genes – PSMB3, CHCHD4, SPDYE6, HSPA5 – were identified for further study based on the novelty. Genes were further chosen based on their involvement in the most enriched pathways seen in our enrichment mapping from data across the whole screen. The final targets identified to study reflect the significant pathways identified in the screen (Figure 2A). All of these targets showed significant depletion in our screen across all four of their guides at both time points (Figure S1D).

**Figure 2:**
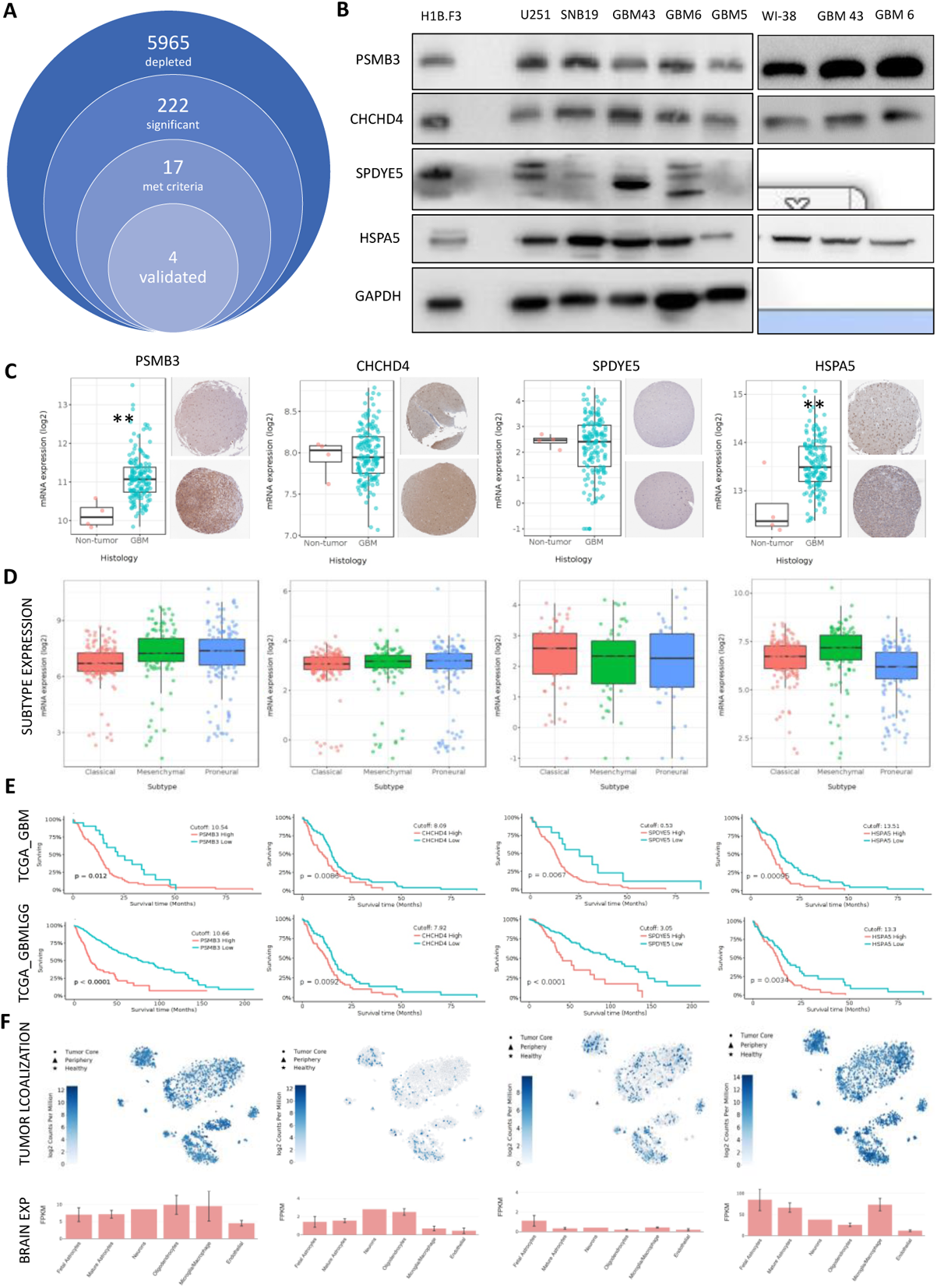
Identified Targets are Associated with Patient Prognosis. (A) Approximately 6000 genes were identified as depleted, of which 222 were significant by multiple algorithms and 17 met criteria of novelty, ability to study, and clinical significance. 4 were chosen for validation to reflect those pathways that were significantly enriched (B) Baseline expression profiles across cell lines used in this study of all genes of interest (C) Expression of targets of interest does not vary by subtype in patient datasets (D) Expression of targets of interest is significantly associated with patient survival in multiple patient datasets (E) Localization of targets within the tumor is not significant for any one location within the tumor. Analysis of subtype expression was performed using Tukey’s Honest Significant Difference (HSD) test and analysis of survival graphs was performed using the log-rank test.

To study these genes, we first performed western blotting to understand the baseline expression of these genes in normal neural stem cells (NSCs) and fibroblasts compared to a range of commercial and patient-derived xenograft (PDX) GBM lines. Corresponding normal cells to cancer lines showed that expression of PSMB3 and HSPA5 is usually elevated compared to normal whereas CHCHD4 and SPDYE5 are not necessarily elevated. This supports the idea that CRISPR screens can powerfully identify genes that are functionally important, which is a departure from the traditional methods of assuming that those genes that are elevated in expression are most important. Notably, our results also reflect a range of expression across different GBM subtypes and different genes, and hence, all experiments were performed in multiple cell lines (Figure 2B). We further validated our results by examining patient datasets and we noted similar trends in expression for the genes of interest. The differences in subtype expression were also reflected in patient datasets. with different subtypes showing variable expression levels for genes of interest (Figure 2D). However, it is essential to note that there are no differences observed under TMZ-based chemotherapy in the expression of these targets. They are targets that act as the tumor is formed, not as a mechanism of resistance to therapy (Figure S2).

Next, we sought to understand the clinical significance of these targets by assessing their relationship to patient survival. Analysis of gene expression compared to survival in multiple patient datasets showed that all four of our targets of interest showed significantly reduced survival with higher expression in patients, reflecting their importance in promoting progression (Figure 2E) [25]. Finally, we assessed localization within the tumor inpatient data at the single-cell level using the publicly available dataset at gbmseq.org [24]. Here, we examined whether there were differences between tumor core and peripheral expression of our targets and noted no significant differences. We further assessed whether certain subpopulations of cells – i.e., vascular, neoplastic, immune, etc. – had significant differences in expression of targets, but we did not note any significant changes in expression based on location within the tumor or subpopulation for any of our targets (Figure 2F). However, we additionally looked at expression in regular brain tissue for our targets, and here we noticed some differences. For example, we noted the SPDYE5 was notably expressed in fetal astrocytes but whereas genes like PSMB3 and HSPA5 were more broadly expressed in all subpopulations (Figure 2F).

### Knockdown of Targets Affects GBM Viability In Vitro & In Vivo

Our screen, at its core, reflected that knocking out these genes should result in reduced viability of cells. To validate that, we first examined correlations between our targets of interest and the well-known marker of proliferation Ki67. Results showed strong positive associations between Ki67 and all four genes of interest, reflecting their importance in proliferation (Figure 3A).

**Figure 3:**
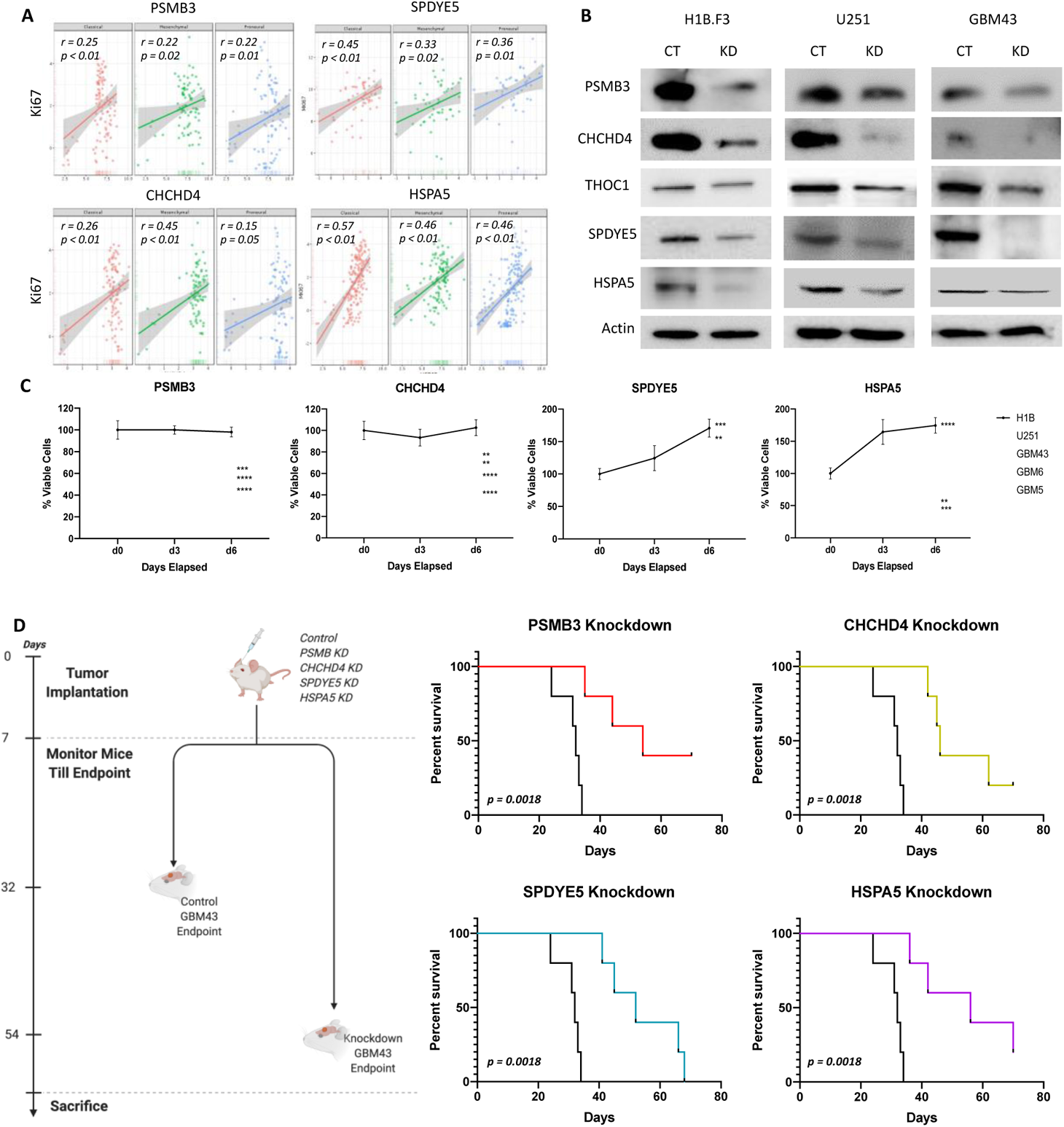
Identified Targets Affect Viability Both In Vitro and In Vivo. (A) Significantly positive correlations were observed between targets of interest and the proliferation marker Ki67 (B) CRISPR-Cas9 based lentiviral knockdowns were established in multiple cell lines and were assessed by western blotting (C) Viability of cells was assessed at multiple timepoints in multiple cell lines, showing that viability was significantly affected by knockdown of target gene compared to control H1B cells (D) GBM43 knockdowns and control cells were implanted in mice per the depicted in vivo scheme. For all targets, knockdown of target genes resulted in significant increases in survival. Analysis was performed in Prism 8, using ANOVA to compare row-means to determine significance or using log-rank tests to determine survival significance *p < 0.05; **p < 0.01; ***p < 0.001; ****p < 0.0001; ns, not significant.

To further validate that knocking out these genes results in reduced cell viability, we developed CRISPR-Cas9 lentiviral knockouts in multiple commercial and PDX GBM lines. Knockdown efficiencies were tested by western blot (Figure 3B). For each gene of interest, three guide plasmids were initially tested in a small set of cell lines, and the most efficient knockout was used to perform experiments across all cell lines (Figure S3A). Cells were then grown for 6 days with regular assessments of cell viability by the MTT assay. Our results showed that, across multiple cancer lines, knocking these genes resulted in significantly reduced viability but that the control NSC line (H1B.F3) remained relatively unaffected (Figure 3C). This result was confirmed visually by microscopic examination (Figure S3B).

Next, we implanted our GBM43 PDX knockout lines in animals and monitored the mice until their endpoint. Across all four genes, there were significant improvements in median survival with the knockout compared to the control. Our control mice had a median survival of approximately 32 days, whereas the PSMB3 knockdown had a median survival of 54 days; the CHCHD4 knockdown had a median survival of 46 days; the SPDYE5 knockdown had a median survival of 52 days, and the HSPA5 knockdown had a median survival of 56 days, all of which were significantly longer than the control (p = 0.0018) (Figure 3D). After mouse death, brains were harvested, and immunofluorescence staining of the tumor tissue was performed to visualize effective and sustained knockdown of these genes (Figure S3C).

### Overexpression of Target Genes Results in Tumorigenesis

We assessed the expression of all four genes in patient tissue samples, comparing cerebral cortex tissues to glioblastoma tissues. Two genes showed consistent upregulations – PSMB3 and HSPA5 – at both the protein and the RNA level (Figure 4A, B) [26]. Therefore, we wanted to investigate whether overexpression of these two genes would result in the transformation of normal cells to a tumorigenic potential.

**Figure 4:**
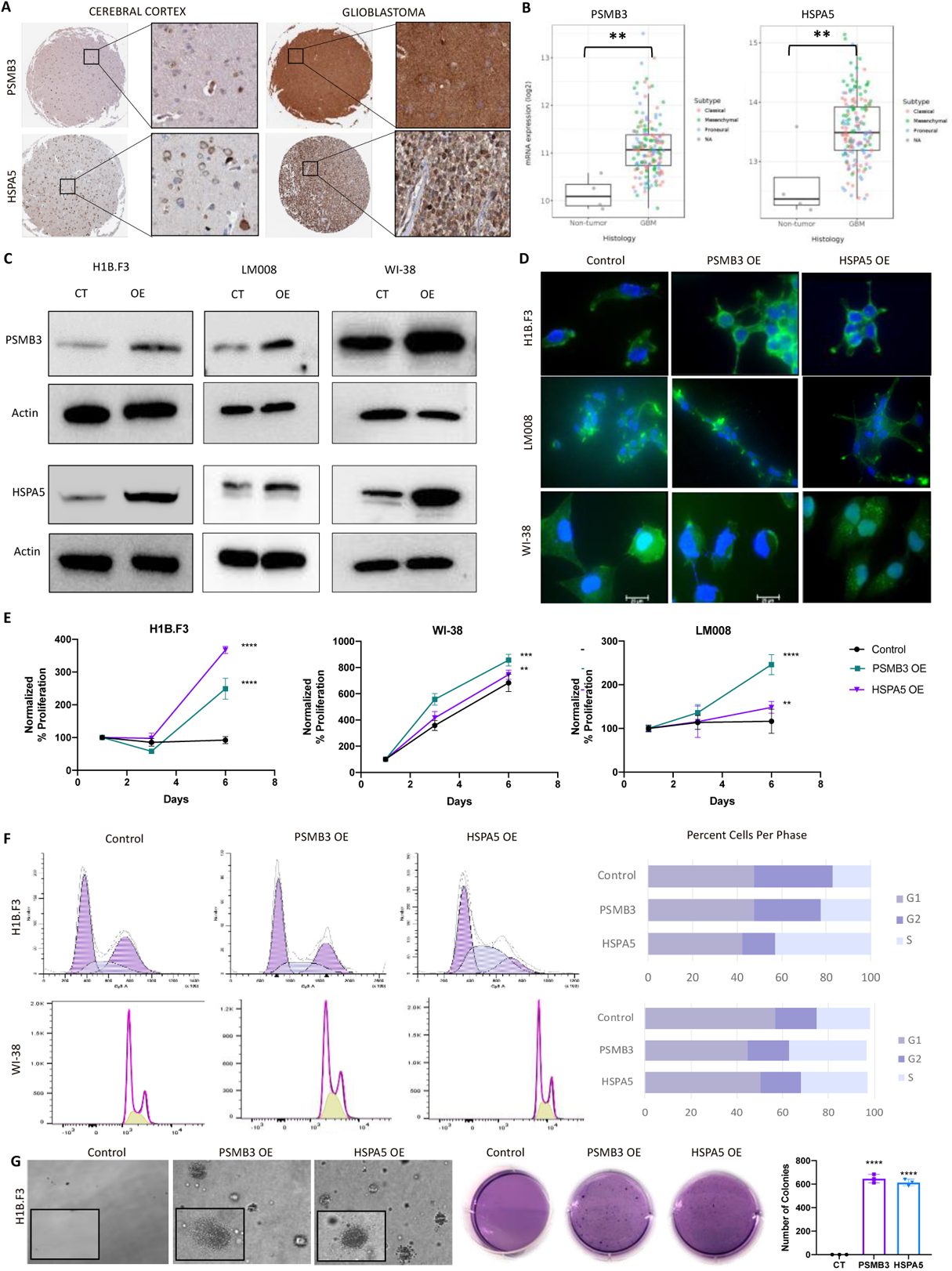
Overexpression of PSMB3 and HSPA5 Results in Tumorigenesis In Vitro. (A) Patient data showed significantly increased expression of PSMB3 and HSPA5 at the protein level (B) Patient data showed significantly increased expression of PSMB3 and HSPA5 at the mRNA level, with no significant differences based on subtype (C) Overexpression lines were established using ORF clones and efficiency of lentiviral transduction was assessed by western blotting (D) Morphology was assessed in each cell line using a phalloidin stain (green) and DAPi (blue). (E) Proliferation of the overexpression lines compared to control was assessed using MTT assays over 1, 3, and 6 days (F) Cell cycle flow cytometry was performed using propidium iodide (PI) staining in overexpression versus control lines (G) Soft agar assays were performed in overexpression versus control and quantified using ImageJ. Analysis was performed in Gliovis using the Tukey’s Honest Significant Test (HSD) or in Prism 8, using ANOVA to compare row-means to determine significance or using log-rank tests to determine survival significance *p < 0.05; **p < 0.01; ***p < 0.001; ****p < 0.0001; ns, not significant.

We overexpressed these genes using ORF clones in three lines of interest – H1B.F3 (NSC), NSC LM008 (NSC), and WI-38 (fibroblast line). The efficiency of overexpression was validated by western blotting and reflected strong overexpression of all genes in our lines of interest (Figure 4C). Morphology of the cells was assessed with a phalloidin stain for actin. Results showed that the overexpression cells seemed to develop longer, spikier shapes than control, potentially reflecting a stronger attachment and a more migratory phenotype (Figure 4D) [27, 28].

Proliferation is a hallmark of tumorigenesis, and thus we assessed it both by MTT assays and by cell cycle flow cytometry. MTTs showed that all three cell lines had significantly increased cell proliferation in the overexpression lines than control (p < 0.001). For example, for H1B.F3 cells, we noted that cells proliferated to 250% and 300% relative to the control, which remained at approximately 100% over our 6-day assay (p < 0.0001). (Figure 4E). This was explained by a significant increase in the percentage of cells found in the S phase in the overexpression cells compared to the control across all three cell types. In H1B.F3 cells, we noted 19% of cells in S-phase as compared to 23% in PSMB3 overexpression and 42% in HSPA5 overexpression (p < 0.001). In fibroblasts, we saw a similar trend, with only 22% of cells in S-phase in the control, relative to 37% in PSMB3 overexpression and 31% in the HSPA5 overexpression (p < 0.001) (Figure 4F).

The gold standard assay for tumorigenesis is the soft agar assay. Tumorigenic cells can grow in this environment despite having no basement membrane to attach to. We performed this assay in multiple cell lines. We showed massive colonies growing after overexpression compared to no colonies in control, providing the most robust evidence that these lines are tumorigenic after overexpression of each gene individually. Specifically, we noted approximately 600 colonies in the PSMB3 and HSPA5 overexpression conditions, relative to no colonies in the control (p < 0.0001) (Figure 4G).

Next, we assessed whether these cells could form tumors in vivo. H1B.F3 cells with control or overexpression lentiviral vectors were injected into mice, and mice were monitored for tumor formation and end point survival. Once the mice had reached their endpoint, brains were harvested and sectioned for histology (Figure 5A). Our results showed extremely rapid deterioration and death of the mice with tumors that had genes overexpressed. PSMB3 overexpression mice had a median survival of 22 days, whereas HSPA OE mice had a median survival of 24 days, whereas all controls remained alive and healthy until an endpoint of 45 days (p = .0039) (Figure 5B). Histological analysis with an in-house licensed neuropathologist showed clear tumor formation in both overexpression mice, representative of a high-grade medulloblastoma. Furthermore, tumors showed strong Ki67 staining, reflecting their highly proliferative nature (Figure 5C).

**Figure 5:**
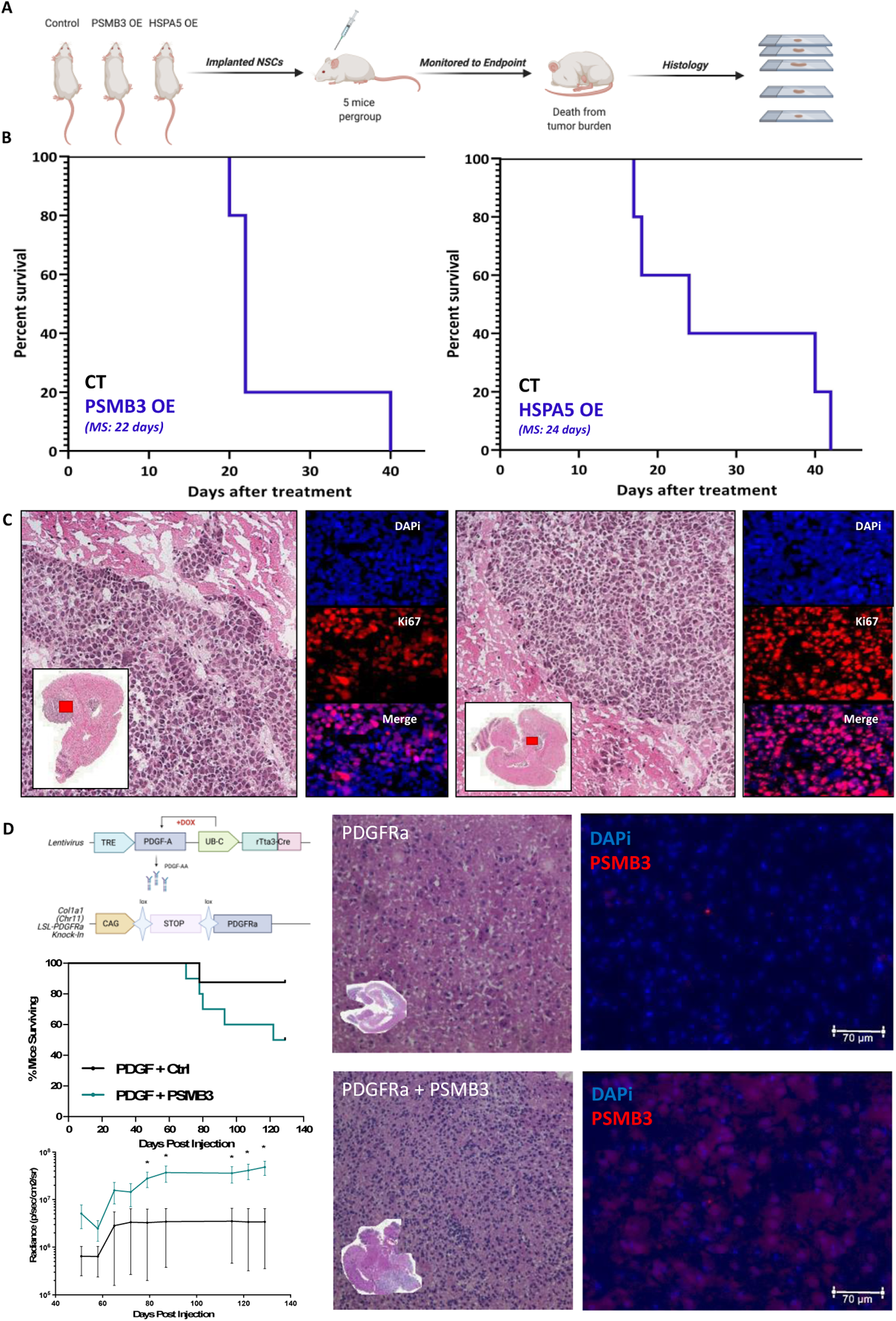
Overexpression of PSMB3 and HSPA5 Results in Tumorigenesis In Vivo. (A) Mice were implanted with control or overexpression NSCs and monitored until the endpoint, after which histology was performed (B) Mice showed significant differences in survival with the overexpression cells as compared to control (C) Histological analysis showed clear tumor formation, with strong Ki67 signaling within the tumor (D) Transgenic mice with PDGFR activation in addition to PSMB3 overexpression formed tumors faster and more aggressively than those without PSMB3 activation Analysis was performed in Prism 8, using ANOVA to compare row-means to determine significance or using log-rank tests to determine survival significance *p < 0.05; **p < 0.01; ***p < 0.001; ****p < 0.0001; ns, not significant.

Furthermore, we performed this experiment in a retroviral inducible transgenic model (Figure 5D). We investigated a previously established PDGFRa-based mouse model for studying glioblastoma in addition to a PDGFRa + PSMB3 overexpression construct [29]. We used a suboptimal dose of virus in order to assess whether tumor formation would still be increased at that lower dose. Results showed that mice died significantly faster with the PSMB3 overexpression vector and that they formed more aggressive, larger tumors by bioluminescence imaging (Figure S4). Overall, these results show that overexpression of these single genes is sufficient to transform a normal NSC line to a tumorigenic phenotype both in vitro and in vivo.

### PSMB3 Knockdown Promotes Proteasome-Mediated Cell Death Through the PSMB3-BIN1-MYC Axis

PSMB3 is a beta subunit of the 19S core proteasome, which plays a key role in the homeostasis of proteins and recycling of amino acids in the cell [13, 30, 31]. Classically, proteins in the cell are ubiquitinated by a chain of ligases once the cell determines they are no longer needed. As this ubiquitin chain grows, the protein is targeted to the proteasome, degrading the protein and recycling free ubiquitin and amino acids for further use in the cell. This process is essential to cellular homeostasis and efficient cell function [13]. It has been implicated as being upregulated in many cancers, and GBM is no exception [11, 12, 14, 16].

Given the importance of the proteasome in cancers and our identification of PSMB3 as a critical target in our screen, we hypothesized that knockdown of PSMB3 would result in dysregulation of the proteasome system and a buildup of ubiquitinated proteins (Figure 6A). We performed a proteasome activity assay, first to assess baseline activity in normal cells versus cancer and showed that GBM cells had almost 300-fold higher proteasome activity than H1B.F3 NSCs (p < 0.001). However, when PSMB3 was knocked down, our data show that proteasome activity diminishes almost 250-fold in cancer, thus reflecting the knockdown of PSMB3 dysregulates the entire proteasome system (p < 0.0001) (Figure 6B). We further showed that the knockdown of PSMB3 in two cancer lines – U251 and GBM43 – showed a buildup of ubiquitinated proteins in the cell, reflecting that the proteasome was no longer functional (Figure 6C). This effect is similar to that of proteasome-targeting drugs. We have shown that treatments with the general proteasome inhibitors bortezomib and MG132 result in downregulation of PSMB3 and a buildup of ubiquitinated proteins (Figure S5).

**Figure 6:**
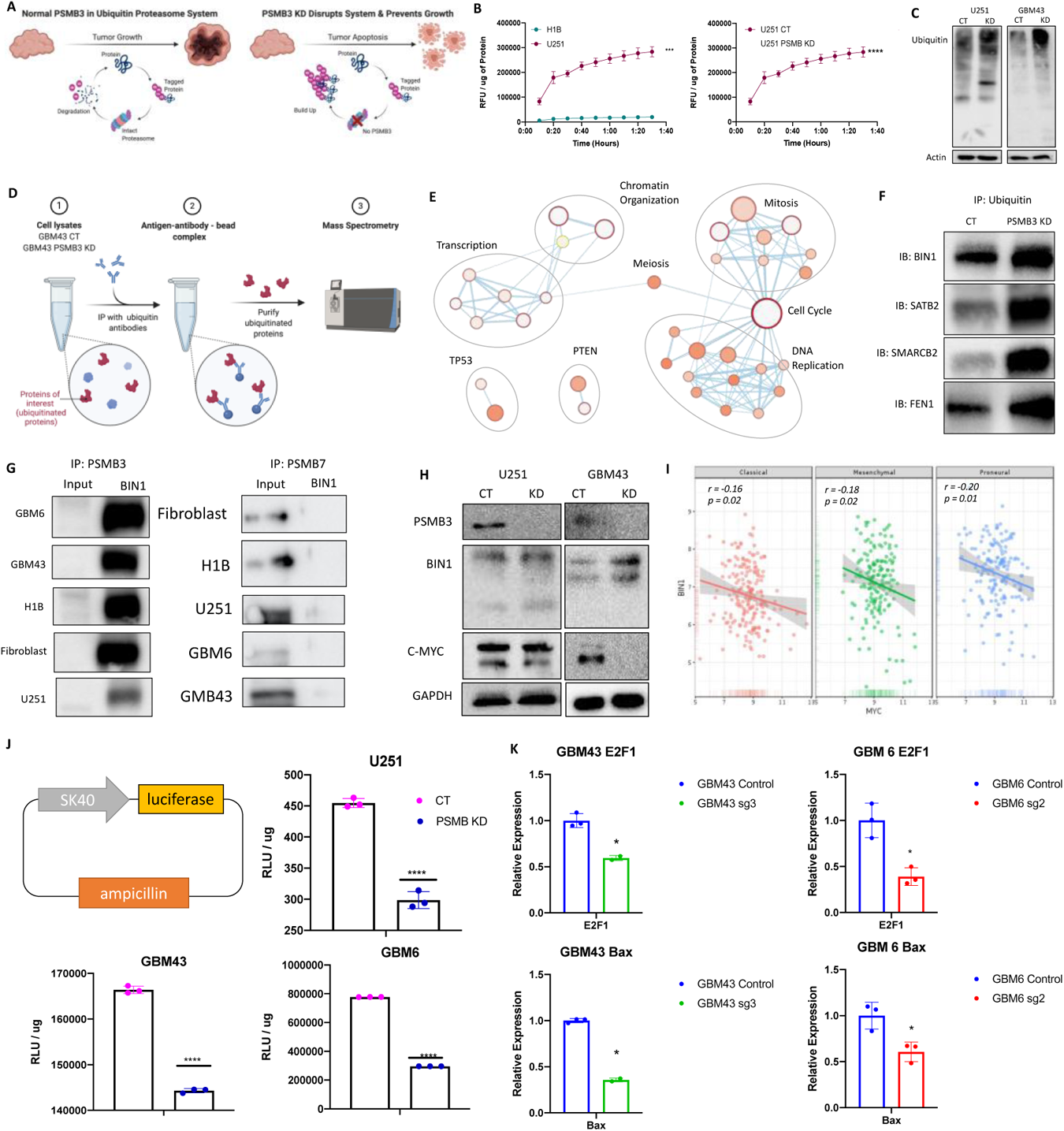
PSMB3 Knockdown Reduces Viability Through the Proteasome-Mediated PSMB3-BIN1-Myc Axis. (A) PSMB3 disrupts the proteasome system and results in buildup of ubiquitinated proteins and apoptosis (B) Comparison of normal versus cancer lines shows that proteasome activity is significantly elevated in cancer compared to normal and knockdown of PSMB3 disrupts proteasome activity (C) Ubiquitinated proteins build up in the PSMB3 knockdown as compared to control in multiple cell lines (D) Immunoprecipitation-mass spectrometry (IP-MS) was performed to assess which proteins are ubiquitinated in cancer (E) Pathways that are ubiquitinated in the knockdown include tumor suppressor pathways and cell cycle and growth pathways (F) Manual validation of multiple targets from the screen reflected ubiquitination increases in the knockdown (G) Examination of specificity of BIN1 binding to PSMB3 revealed that BIN1 is highly specific to PSMB3 (H) Western blot shows that PSMB3 influences BIN1 activity which then influences c-myc activity in the cell (I) BIN1 and c-myc expression are inversely correlated in patient datasets (J) Myc activity is downregulated with the PSMB3 knockdown as measured by a luciferase reporter (K) Myc downstream genes E2F1 and BAX are transcriptionally downregulated in the PSMB3 knockdown. Analysis was performed in Prism 8, using ANOVA to compare row-means to determine significance or using log-rank tests to determine survival significance *p < 0.05; **p < 0.01; ***p < 0.001; ****p < 0.0001; ns, not significant.

Based on our observation that PSMB3 is associated with cell viability and that PSMB3 knockdowns disrupt the proteasome system, our next step was to identify what pathways are being targeted by the proteasome in cancer. To do this, we performed immunoprecipitation mass spectrometry (IP-MS) with antibody against ubiquitin in GBM43 control versus a PSMB knockdown (Figure 6D). Results identified 62 proteins that were highly ubiquitinated in the knockdown. This represents those proteins being targeted by ubiquitin in a cancer cell that can no longer degrade the proteins due to proteasome dysregulation. Enrichment analysis of the proteins showed vital tumor suppressor pathways such as TP53 and PTEN, as well as a number of growth processes that require regular recycling of necessary proteins (e.g., cell cycle, mitosis, replication, etc.) (Figure 6E). Results of the IP-MS experiment were validated by immunoprecipitation-western blotting (IP-WB). Here, we show four of the identified targets. - BIN1, SATB2, SMARCB2, and FEN1 – all with significantly increased ubiquitination in the knockdown compared to the control (Figure 6F).

BIN1 was of particular interest as it is known to act as a tumor suppressor by downregulating c-myc [32–34]. Therefore, targeting BIN1 by PSMB3 within the proteasome could be part of the mechanism by which cancer cells survive. To further investigate this axis, we first identified whether or not BIN1 was PSMB3-specific. We first performed IP-WB investigating BIN1 binding to PSMB3 versus PSMB7, another subunit of proteasome. Results showed that BIN1 bound highly specifically to PSMB3 (Figure 6G). Investigation of this PSMB3-BIN1-myc axis in two cancer lines – U251 and GBM43 – showed that PSMB3 knockdown was associated with BIN1 elevations and c-myc downregulation in cancer cells, reflecting that this may be a reason why the PSMB3 knockdown is toxic to cells (Figure 6H). This was further evidenced by a negative correlation between BIN1 and c-myc expression observed in patient samples across all tumor subtypes (Figure 6I). Furthermore, a myc activity reporter assay performed in multiple cancer lines reflected a similar trend of myc activity downregulation with PSMB3 knockdown (Figure 6J). Examination of transcriptionally activated myc downstream targets by qPCR further supported this axis, showing that both E2F1 and BAX, known targets of myc, had reduced transcription at the mRNA level after PSMB3 knockdown [35–37] (Figure 6K).

Conversely, PSMB3 upregulation is associated with increases in myc expression, seen both in patient level RNA data and in our single-cell data. We further validated this in our PSMB3 overexpression lines by western blotting and with a luciferase-based myc reporter assay. Together, this further reflects the important role of PSMB3 in modulating myc expression and thus affecting tumor progression (Figure S6).

### PSMB3 Overexpression Promotes Tumorigenesis Through Notch Activation

Our next step, given that the knockdown of PSMB3 interferes with the proteasomal function, was to assess the effect of PSMB3 overexpression on proteasomal function. Interestingly, we found that overexpression of PSMB3 did not significantly alter proteasome activity (Figure 7A). Given this, we determined that it is possible that in the pre-transformation condition PSMB3 may contribute to oncogenesis in a proteasome-independent alternative pathway.

**Figure 7:**
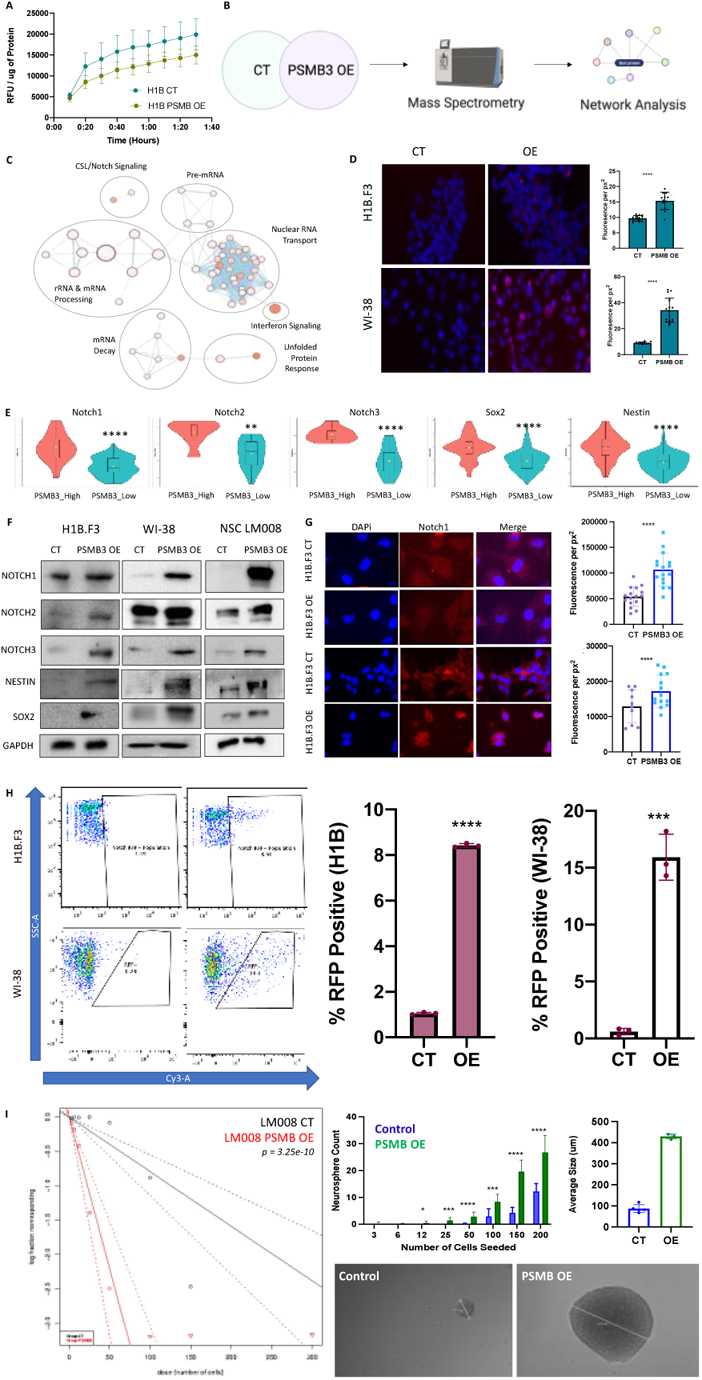
PSMB3 Overexpression Acts Outside the Proteasome to Activate Notch and Stemness Genes. (A) Proteasome activity is not significantly altered in H1B.F3 cells with PSMB3 overexpression, suggesting a non-proteasomal function is driving tumorigenesis in the PSMB3 overexpression (B) H1B.F3 cells were transduced with control or overexpression lentivirus and were sent for proteomics (C) Enrichment mapping showed enrichment of Notch pathways and growth and transcription pathways (D) Immunohistochemistry in patient samples shows that there is a higher level of nuclear localization of PSMB3 in more aggressive tumors (E) Single cell analysis of primary human gliomas implanted in mice reflects that higher PSMB3 expression is associated with higher Notch and stemness marker expression (F) Analysis of expression of notch isoforms and stemness markers by western blot reflected that these markers were elevated in the overexpression as compared to control (G) Notch1 is elevated in the nucleus of PSMB3 overexpression cells shown by immunofluorescence (H) Overexpression of PSMB3 results activation of Notch1 downstream markers, as evidenced by a RFP-based promoter-reporter assay (I) PSMB3 overexpression results in functional stemness, as evidenced by a neurosphere assay in multiple cell lines. Analysis was performed in Prism 8, using ANOVA to compare row-means to determine significance or using log-rank tests to determine survival significance *p < 0.05; **p < 0.01; ***p < 0.001; ****p < 0.0001; ns, not significant.

To further investigate the non-proteasomal function, we performed proteomics on H1B.F3 control cells versus the PSMB3 overexpression cells, and then examined network enrichment of those genes that were elevated in the overexpression (**Figure 7B**). Analysis reflected significant enrichment of tumorigenic pathways such as Notch signaling pathways in addition to nuclear transport and rRNA and mRNA processing (**Figure 7C**). Taken together, this suggests that PSMB3 may be translocating to the nucleus when overexpressed in addition to being involved in activating notch signaling pathways and thus promoting tumorigenesis. Although this is an area with a paucity of studies, a few other studies have shown that isolated proteasome subunits may act as transcription factors in the nucleus, supporting our hypothesis [38–40].

To validate that PSMB3 is acting in the nucleus at higher levels of expression, we first performed immunohistochemistry in patient samples and were able to show that PSMB3 preferentially localizes to the nucleus at higher expression scores. We additionally performed immunocytochemistry and noticed increased PSMB3 localization to the nucleus in our overexpression cell lines (**Figure 7D, S6**). Next, we examined notch pathway activation at the single-cell level in primary GBM in an *in vivo* mouse model. Cells were stratified based on whether they had low or high expression of PSMB3, and each group was examined for expression of notch isoforms and stem cell markers. Results reflected a similar trend, showing that higher PSMB3 expression was associated with higher activation at the mRNA level of notch isoforms and stem cell markers (**Figure 7E**). This was additionally supported at the protein level. Through western blotting in multiple normal cell lines, we found that overexpression of PSMB3 resulted in overexpression of multiple Notch isoforms in addition to classically known cancer stem cell markers Sox2 and Nestin (**Figure 7F**) [41–43]. This is further bolstered by the fact that PSMB3 knockdown in cancer does result in alterations of the Notch pathway and stemness markers, as shown at the RNA level in patient datasets and as verified by western blotting in our glioblastoma PSMB3 knockdown cell lines (Figure S7).

To understand how Notch protein upregulation was influencing the cells, we examined the localization of Notch1 in the cells. Immunofluorescent examination of Notch1 showed that there was an elevation of Notch1 protein expression in the nucleus in PSMB3 overexpression cells (**Figure 7G**). This suggests that once transcribed, Notch1 preferentially enters the nucleus to promote transcription of downstream genes to promote a cancer phenotype. To assess whether Notch1-promoted transcription is occurring in the cells, we performed a Notch1 reporter assay with an RFP-linked promoter-reporter system. Results in multiple cell lines showed that in the overexpression condition, there is significantly more activation of Notch downstream targets as compared to control (*p* < 0.0001) (**Figure 7H**). Finally, to perform a functional assay of the stemness phenotype we see upregulated here, we performed an extreme limiting dilution neurosphere assay in multiple cell lines. We found that overexpression of PSMB3 significantly increased sphere formation capacity and thus functional stemness and tumorigenicity of the cells (**Figure 7I, S8**) Together, these results clearly reflect that PSMB3 activates Notch and stemness pathways, which then have a functional tumorigenic impact on the cell population.

## Discussion

Glioblastoma is a debilitating and devastating disease with a terrible prognosis in patients. As such, it is a disease that desperately requires innovative approaches in order to identify actionable targets. Previously, it has been shown that CRISPR-Cas9 whole-genome screens can be valuable tools for identifying previously unknown genetic vulnerabilities in cell population in an unbiased manner [8, 9]. However, this technique has not yet been applied to understand novel therapeutic targets in GBM, representing untapped promise as these screens are an extremely valuable tool for us to approach the genome and have an unbiased view of what is functionally important for the proliferation and aggressiveness noted consistently in glioblastoma.

Here, we perform a whole-genome CRISPR-Cas9 screen to identify almost 6000 genes that are important for viability and proliferation of GBM cells. Of these, approximately 220 were significant across multiple analysis methods and multiple timepoints, all of which represent highly promising targets for combating GBM. Furthermore, we were able to identify key cellular programs or pathways driving this process, further defining which pathways are important to target in the future. Validation of four of these targets, all previously unstudied, shows highly promising results with all four showing significant effects on GBM viability *in vitro, in vivo,* and in various patient datasets. These results bolster the accuracy of the screen and validate that the targets we have identified universally have a role in cell proliferation. Although we validated four of these targets, the fact that all four targets demonstrated efficacy suggests that the 200+ other targets identified may additionally hold promise for future therapeutics.

Two of these targets are known to consistently have elevated expression in glioblastoma. To understand whether they can contribute to tumorigenesis, we established lentiviral overexpression cell lines in two neural stem cell lines and in one non-neural line, lung-derived fibroblasts. Our goal in using fibroblasts was to understand whether the effects of these genes could extend beyond neural cells and play a contributory role in other cancers outside the brain. Our results showed that overexpression of these targets in normal cell lines, both in neural stem cells and fibroblasts, alters morphology, increases proliferation, and results in growth of colonies on soft agar, all hallmarks of tumorigenesis. Furthermore, we found that these cells were able to form tumors *in vivo* and resulted in significantly accelerated fatality in mice. Together, these results are startling and demonstrate additional importance of these genes not initially suggested by our CRISPR-Cas9 screen. This means that these genes, in isolation, when overexpressed, contribute significantly to tumorigenesis, not just in neural lines and neural cancers but potentially in other cancers also.

PSMB3 was of particular interest to us throughout this study as it is a core subunit of the beta ring of the 19S proteasome. The proteasome has a long history of being studied in cancer and it is well-known to be upregulated in most human cancers [14–16]. Proteasome inhibitors have been attempted and have had clinical success in other cancers. However, there are notable gaps in the literature around the mechanism of these proteasome inhibitors in any cancer, including glioblastoma. In order to understand why the proteasome is important in glioblastoma and to understand how it may be effectively targeted, it is essential for us to understand the mechanism by which it promotes tumorigenesis. Here, we show that the proteasome in glioblastoma specifically targets tumor suppressor pathways in addition to a range of growth proteins that are constantly recycled in any actively growing cell. This sheds important light on why the proteasome plays such a key role in cancers and provides additional points of inhibition along the proteasome pathway to allow synergistic therapies to be more effective.

Another aspect of PSMB3 that remains extremely understudied is potential non-proteasomal functions of the subunit. This is an area of new interest in the literature but there remains a dearth of studies that focus on this topic [38, 40]. Although many proteasomal subunits appeared in our screen, this one was by far the most significantly depleted. Thus, it must have additional importance outside of its role in the proteasome. It is conceivable that pre-transformation, PSMB3 has additional roles that non-proteasomal. This is certainly what our data suggests as overexpression of PSMB3 did not significantly alter proteasome activity. Our data shows that PSMB3 translocates to the nucleus, especially in more aggressive cancers, and likely plays roles in transcription and activation of Notch isoforms and the stemness program within cells.Individual targeting of the proteasome subunits rather than generalized targeting of the proteasome as a whole may be a viable strategy in the future. In addition, various other subunits may have important functions that remain unidentified in the literature currently. We believe this understanding of the proteasome and non-proteasomal functions of PSMB3 in glioblastoma helps shed light on one more aspect of what makes this tumor so deadly and provides a novel avenue by which to prevent progression of primary glioblastoma.

## Supplementary Figures

**Figure S1:**
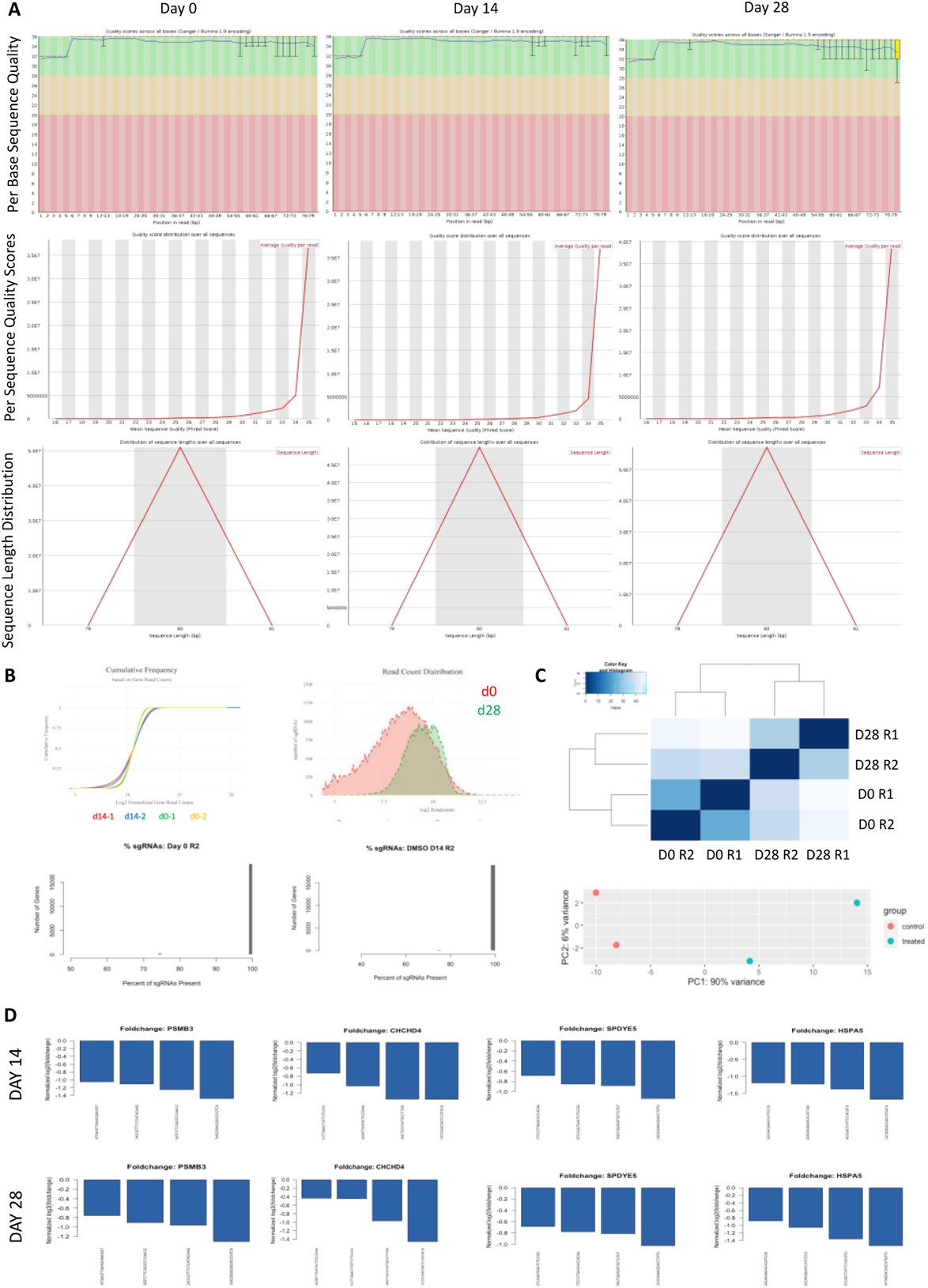
CRISPR-Cas9 Screen Quality. (A) Quality of sequencing data is appropriate for all time points (B) Cumulative frequency at days 0 and 28 was assessed and reflected an appropriate distribution. Guide coverage is also appropriate across our screen with limited guide loss over time (C) Replicates show appropriate similarity to each other in our screen for day 0 and d28 by distance analysis and by PCA (D) All chosen targets show significant depletion of all four guides at both timepoints of our screen.

**Figure S2:**
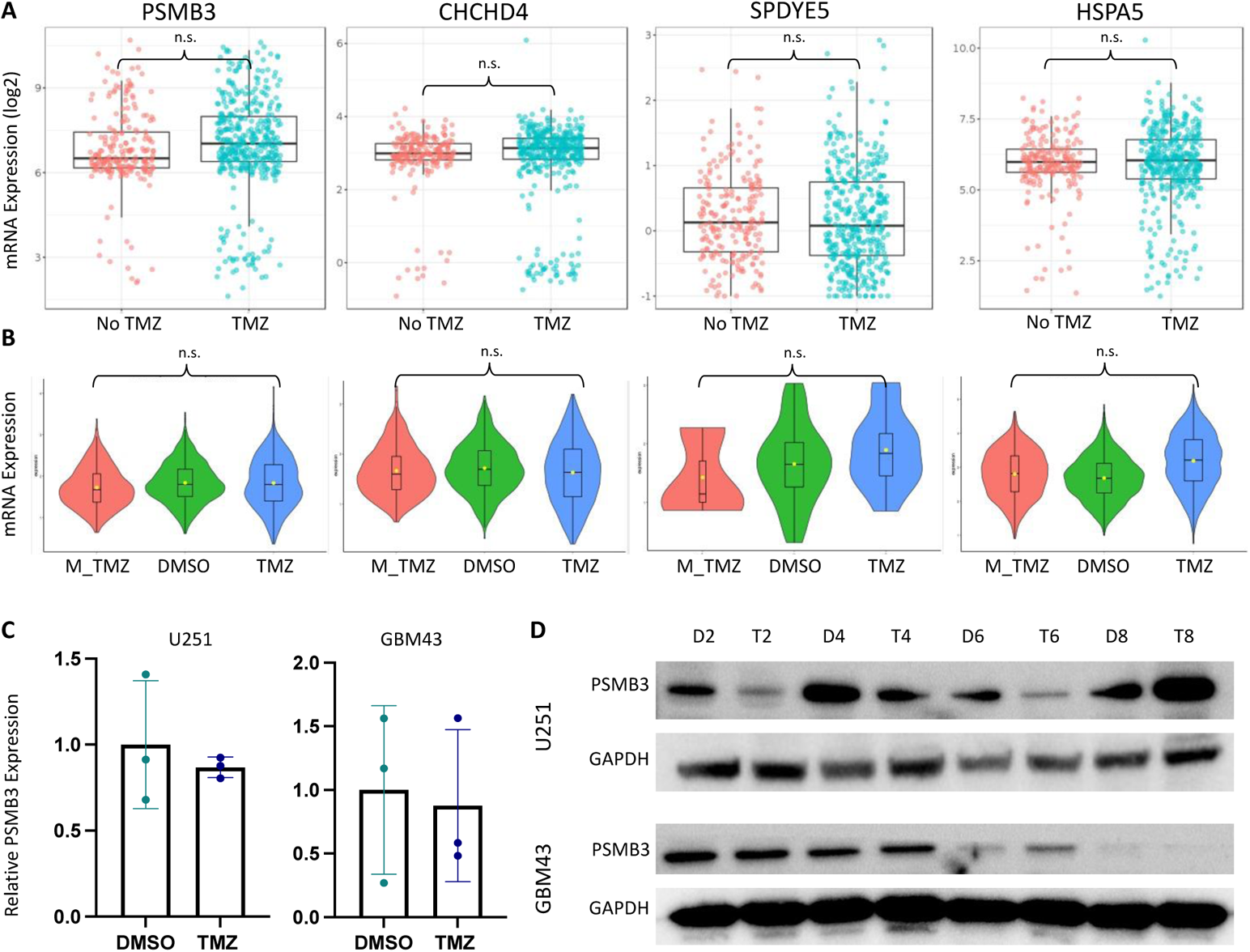
TMZ-Based Chemotherapy Does Not Impact Targets. (A) Patient-level RNA-Seq data shows no significant changes between patients who received chemotherapy and those who did not in target gene expression (B) Single cell data from primary gliomas implanted in a mouse model show no difference in expression of targets before, during, or after therapy (C) qPCR analysis of PSMB3 expression in cancer lines treated with TMZ shows no change in PSMB3 expression at two days of treatment (D) Western blotting over eight days of treatment with TMZ shows no significant change or pattern in expression of PSMB3 with TMZ therapy. Analysis was performed in Prism 8, using ANOVA to compare row-means to determine significance or using log-rank tests to determine survival significance *p < 0.05; **p < 0.01; ***p < 0.001; ****p < 0.0001; ns, not significant.

**Figure S3:**
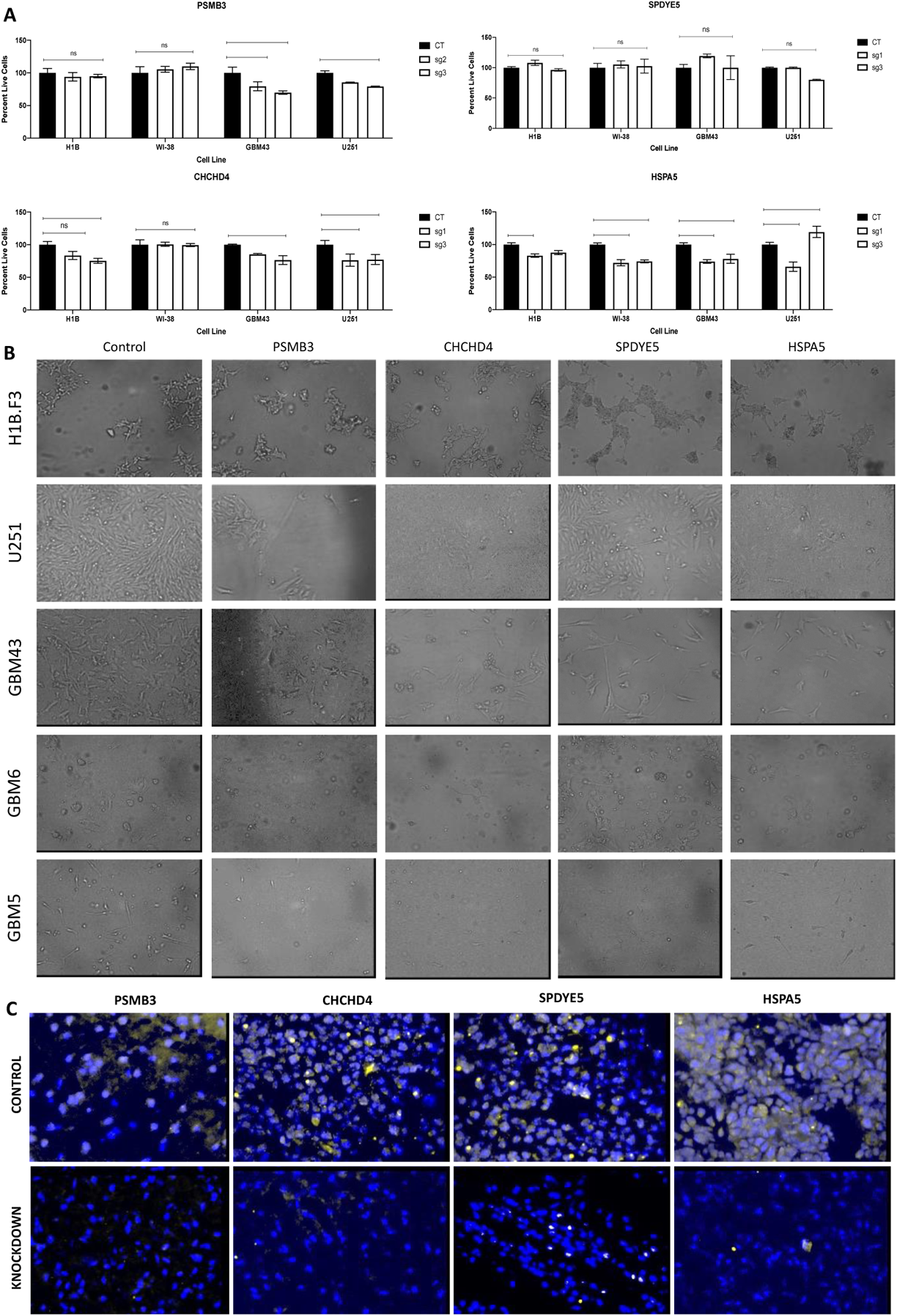
Knockdown of Targets Affects Viability in Multiple Cell Lines. (A) Use of multiple guide RNAs to knockdown targets of interest in four cell lines demonstrates significant effects on viability (B) Cell viability images in a single guide RNA knockdown for each cell line and target of interest (C) Histology from in vivo knockout experiment demonstrates that the knockouts of targets of interest were sustained. Analysis was performed in Prism 8, using ANOVA to compare row-means to determine significance or using log-rank tests to determine survival significance *p < 0.05; **p < 0.01; ***p < 0.001; ****p < 0.0001; ns, not significant.

**Figure S4:**
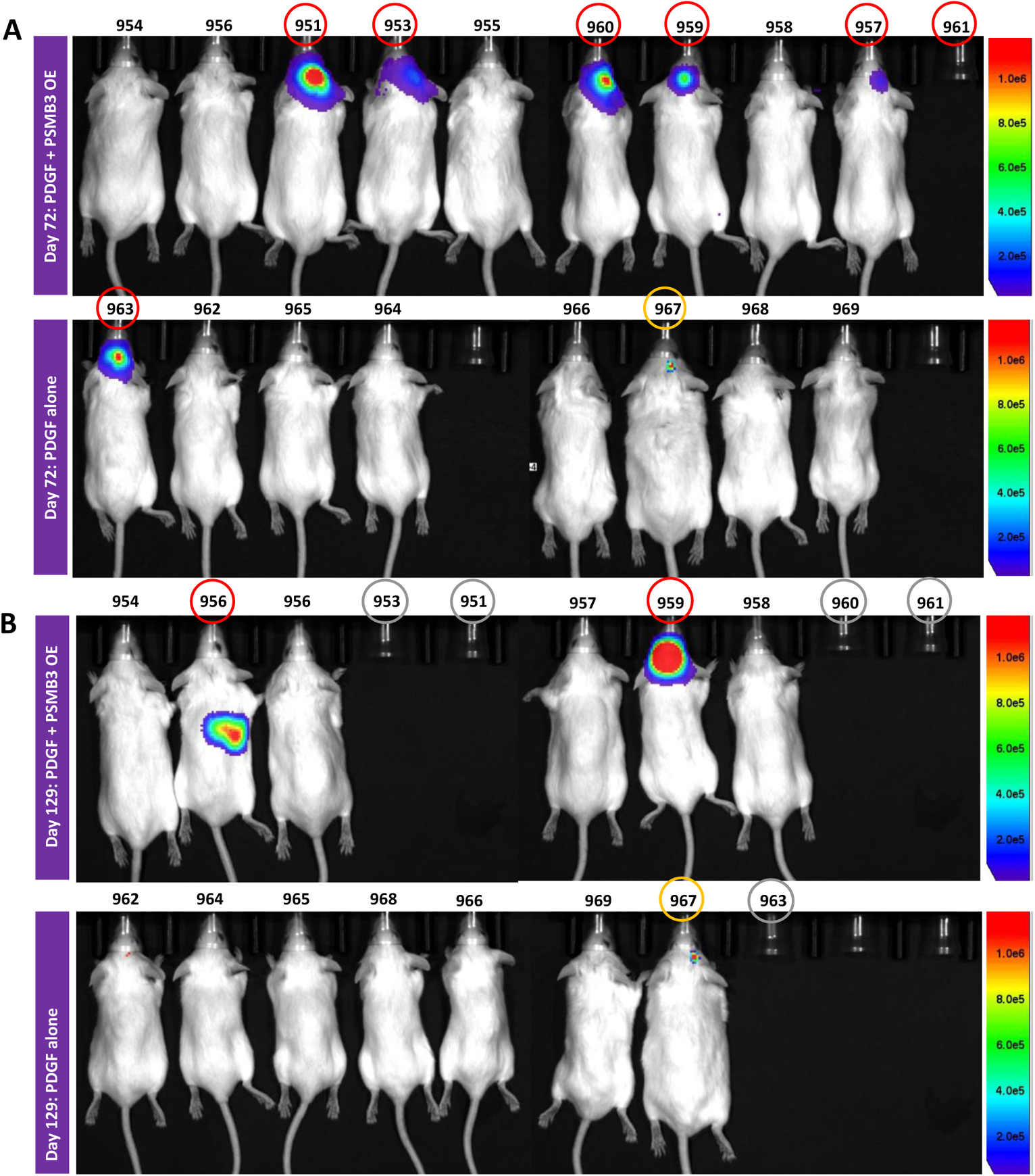
PSMB3 Overexpression in Transgenic Mice Results in Aggressive Tumors. (A) Transgenic mice were assessed by BLI and are depicted here 72 days following injection and (B) 129 days following injection

**Figure S5:**
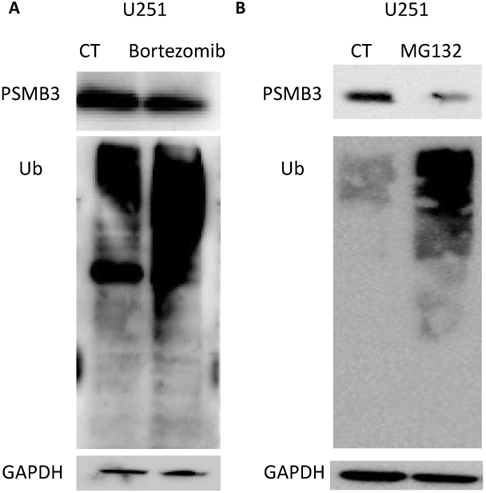
Treatment of Cells with Proteasome Inhibitors Affects PSMB3 Expression. (A) Treatment of cells with the clinical proteasome inhibitor bortezomib downregulates PSMB3 expression (B) Treatment of cells with the generic proteasome inhibitor MG132 downregulates PSMB3 expression.

**Figure S6.**
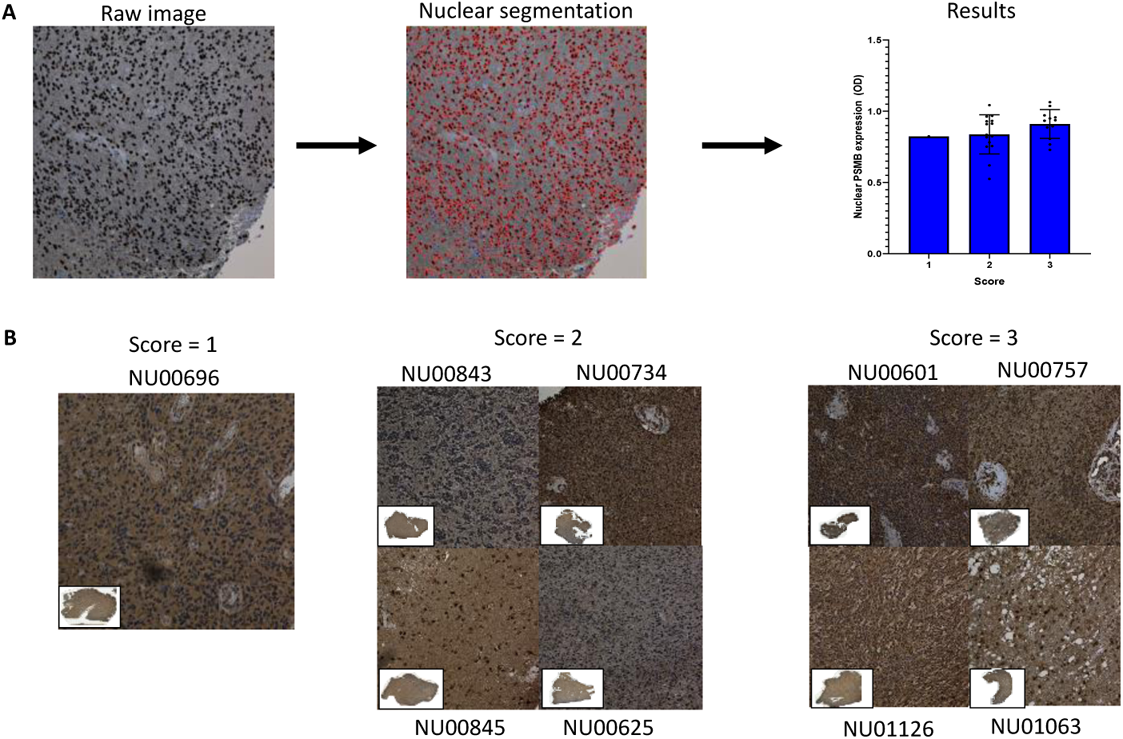
Patient Histology Shows Elevated Nuclear Localization of PSMB3. (A) Nuclear localization of PSMB3 was assessed using the HistoQuest program (B) Raw histological images used for analysis of data

**Figure S7:**
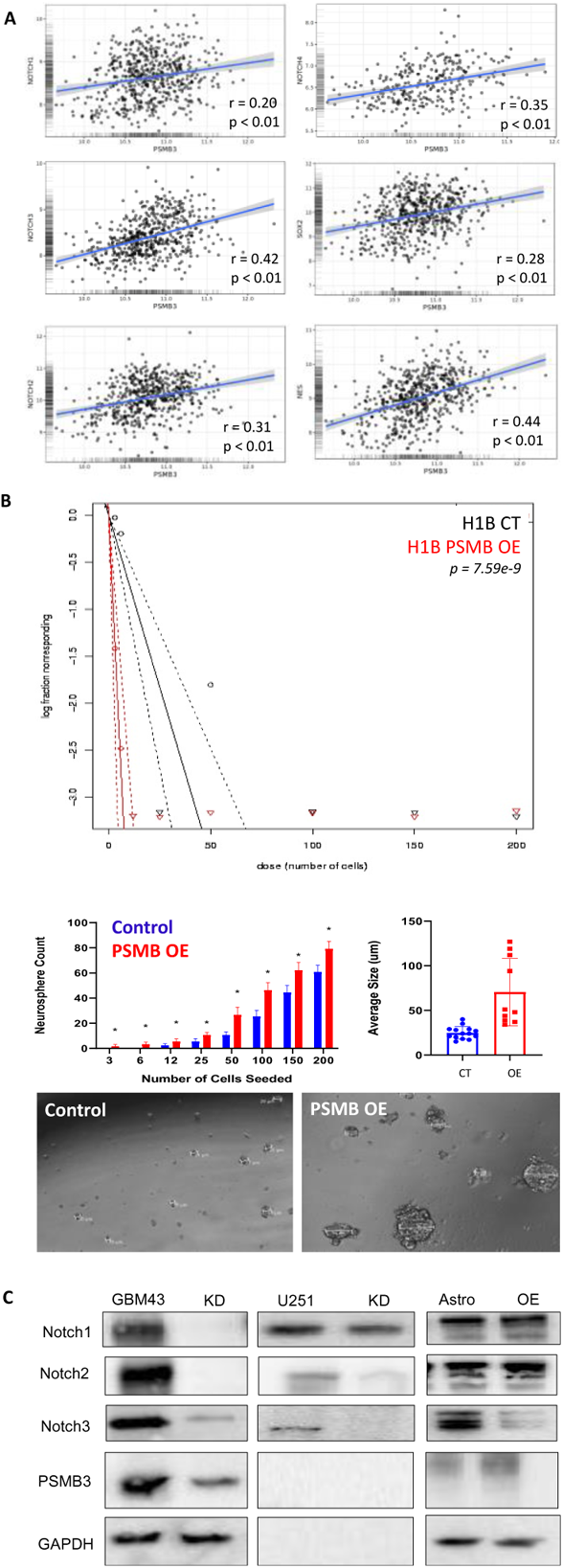
PSMB3 Knockdown Downregulates Notch Isoforms and Stemness Markers. (A) PSMB3 expression is associated with Notch isoform and stemness marker expression (B) H1B PSMB3 overexpression demonstrates increased functional stemness, as evidenced by a neurosphere assay, both by the quantitiy and the size of spheres formed (C) U251 and GBM43 cells were transduced with PSMB3 knockdown vectors and blotted for notch isoforms and stemness markers. Both cell lines universally show downregulation of these markers with PSMB3 knockdown. Interestingly, astrocytes do not transform in the same way as neural stem cells do and continue to show stemness downregulation with the PSMB3 overxpression Correlation analysis was performed using Pearson’s Correlation Coefficient to determine significance *p < 0.05; **p < 0.01; ***p < 0.001; ****p < 0.0001; ns, not significant.

## Acknowledgements

This work was supported by the National Institute of Neurological Disorders and Stroke grant 1R01NS096376, 1R01NS112856 the American Cancer Society grant RSG-16-034-01-DDC (to A.U.A.) and P50CA221747 SPORE for Translational Approaches to Brain Cancer.

The results published here are in part based upon data generated by the TCGA Research Network: https://www.cancer.gov/tcga, and were further analyzed through GlioVis. In addition, these results use data generated by the Human Protein Atlas and GBMSeq (Gephart Lab, www.gbmseq.org). Figures, in part, were generated using BioRender (www.biorender.com).

## Author Contributions

Conceptualization: Shivani Baisiwala, Adam M Sonabend, Atique U Ahmed Methodology: Shivani Baisiwala, Shreya Budhiraja, Adam M Sonabend, Atique U Ahmed

Validation, Formal Analysis, Investigation: Shivani Baisiwala, Shreya Budhiraja, Li Chen, Andrew J Zolp, Ella N Perrault, Khizar Nandoliya, Crismita Dmello, Cheol H Park, Miranda R Saathoff, Gabriel Dara, Jack M Shireman, Peiyu Lin, Katy McCortney

Resources: Atique U Ahmed, Adam M Sonabend, Craig Horbinski Data Curation & Draft Preparation: Shivani Baisiwala

Review & Editing: Shivani Baisiwala, Atique U Ahmed, Shreya Budhiraja, Khizar Nandoliya, Adam M Sonabend

Supervision, Project Administration, Funding Acquisition: Atique U Ahmed, Adam M Sonabend

